# *Acvr1b* loss increases formation of pancreatic precancerous lesions from acinar and ductal cells of origin

**DOI:** 10.1101/2023.06.08.544226

**Authors:** Kiyoshi Saeki, Ian S. Wood, Wei Chuan K. Wang, Shilpa Patil, Yanping Sun, David F. Schaeffer, Gloria H. Su, Janel L. Kopp

## Abstract

**Background & Aims:** Pancreatic ductal adenocarcinoma (PDAC) can develop from precursor lesions, including pancreatic intraepithelial neoplasia (PanIN) and intraductal papillary mucinous neoplasm (IPMN). Previous studies indicated that loss of *Acvr1b* accelerates the Kras-mediated development of papillary IPMN in the mouse pancreas, however, the cell type predominantly affected by these genetic changes remains unclear.

**Methods:** We investigated the contribution of cellular origin by inducing IPMN associated mutations-KRAS^G12D^ expression and *Acvr1b* loss - specifically in acinar (*Ptf1a^CreER^;Kras^LSL-G12D^;Acvr1b^fl/fl^*mice) or ductal (*Sox9CreER;Kras^LSL-G12D^;Acvr1b^fl/fl^*mice) cells in mice. We then performed MRI imaging and a thorough histopathological analysis of their pancreatic tissues.

**Results:** The loss of *Acvr1b* increased the development of PanIN and IPMN-like lesions when either acinar and ductal cells expressed a Kras mutation. MRI, immunohistochemistry and histology revealed large IPMN-like lesions in these mice that exhibited features of flat, gastric epithelium. In addition, cyst formation in both mouse models was accompanied by chronic pancreatitis. Experimental acute pancreatitis accelerated the development of large mucinous cysts and PanIN when acinar, but not ductal, cells expressed mutant Kras and lost *Acvr1b*.

**Conclusion:** These findings indicate that loss of *Acvr1b* in the presence of the Kras oncogene promotes the development of large and small precancerous lesions from both ductal and acinar cells. However, the IPMN-like phenotype was not equivalent to that observed when these mutations were made in all pancreatic cells during development. Our study underscores the significance of the cellular context in the initiation and progression of precursor lesions from exocrine cells.

## Introduction

Pancreatic cancer initiation is associated with three main types of precancerous lesions – Pancreatic Intraepithelial Neoplasia (PanIN), Intraductal Papillary Mucinous Neoplasm (IPMN) and Mucinous Cystic Neoplasm (MCN). PanIN and IPMN lesions are both mucinous and clinically they are largely distinguished by their size, with IPMN being a larger lesion type that can be detected by magnetic resonance imaging (MRI). Adult pancreatic ductal and acinar cells can both give rise to precancerous lesions in animal models^1–5^ when mutations in *Kras*, the most commonly altered oncogene in pancreatic cancer, are present. However, the majority of gene mutations and pathway alterations associated with pancreatic cancer have been studied in the context of pan-pancreatic genetic changes induced during pancreatogenesis.^6^ Thus, how a particular gene affects the initiation and development of precancerous lesions and pancreatic cancer from either acinar or especially ductal cells is largely unknown.

Activin A receptor type 1B (ACVR1B), is a member of the transforming growth factor - beta (TGF-β) superfamily of receptor serine/threonine kinase. Compared to the TGF-β and BMP pathways within the TGF-β superfamily, our understanding of activin signaling is not as comprehensive. Activins exert their effect by binding to either ACVR2A or ACVR2B (Type II receptors), which prompts the recruitment of a type I receptor, like ACVR1B. The activated receptor complex then activates SMAD2 and SMAD3, which forms a complex with SMAD4 and regulates gene transcription in the nucleus. Activin signaling plays a crucial role in several cellular processes, such as cell proliferation,^7–9^ cell cycle arrest,^10^ and apoptosis.^11,12^ Dysregulation of activin signaling components has been implicated in several diseases, including cancer. Biallelic inactivation of *ACVR1B* has been reported in pancreatic cancer.^13,14^ Mutations in *SMAD2/3/4*, which are downstream mediators of activin/TGFβ signaling, are also found in many cancers,^15^ with 55% of pancreatic cancers having *SMAD4* loss.^16^ Loss of *SMAD4* was primarily found in PDAC, but not the precancerous lesions. However, both *Smad4* loss and *Acvr1b* loss in the pancreas using animal models is associated with the greater development of PanIN lesions, cystic pancreatic precursors, and pancreatic cancer development in the context of oncogenic Kras,^17–20^ suggesting that the Activin arm of TGFβ superfamily can affect the initiation and development of precursor lesions.

In our previous studies, we showed that loss of *Acvr1b* in mouse pancreas using a mosaic *Pdx1Cre*^21^ allele led to chronic pancreatitis around 9 months of age.^20^ When these alleles were combined with the oncogenic *Kras^LSL-G12D^* allele,^21^ these mice developed cystic gastric and pancreatobiliary IPMN-like lesions beginning around 34 days of age, as well as numerous PanIN and pancreatic cancer.^20^ These data suggested that loss of *Acvr1b* in the context of oncogenic Kras resulted in the induction of IPMN-like lesions. In *Pdx1Cre;Kras^LSL-G12D^;Acvr1b^fl/fl^* (AKpdx1) mice gene mutations are targeted to early pancreatic progenitor cells, resulting in the genetic changes occurring in the developing endocrine, acinar, and ductal cell compartments. This made it unclear which specific cell types were contributing to the development of the IPMN-like lesions in AKpdx1 mice. To determine the consequence of *Acvr1b* loss specifically in acinar or ductal cells in the pancreas in the context of oncogenic Kras, we crossed *Kras^LSL-G12D^;Acvr1b^fl/fl^*mice to mice expressing the CreER protein in an acinar- or ductal-cell-specific manner and examined the development of lesions by MRI and histopathology. We found that *Acvr1b* loss and activation of *Kras* in either the acinar or ductal cells results in chronic pancreatitis, increased formation of MRI-detectable, mucinous cysts, and increased microscopic precancerous lesions. Although some of these large cysts were mucinous, only AKpdx1 mice formed complex papillary IPMN-like lesions. This suggests that the phenotype in AKpdx1 mice is potentially the consequence of a complex interplay of events occurring during development of the pancreas in the context of these mutations or a result of mutations being present in large numbers of acinar and ductal cells.

## Results

### *Acvr1b* loss in Kras^G12D^-expressing acinar or ductal cells promotes cyst formation

To determine the phenotypic consequences of *Acvr1b* loss in the context of Kras^G12D^ expression in postnatal ductal or acinar cells, we generated mouse models in which these genetic changes were induced in an acinar-cell-type- or ductal-cell-type-specific manner. Specifically, we used the tamoxifen-inducible *Ptf1a^CreER^* allele^22^ to induce Cre-dependent recombination of the *Kras^LSL-G12D^* ^21^ and *Acvr1b^flox^* ^20^ conditional alleles to express Kras^G12D^ and ablate *Acvr1b* expression (Fig. 1A) in juvenile-aged acinar cells. By combining all of these alleles through cross-breeding, we generated *Ptf1a^CreER^;Kras^LSL-G12D^;Acvr1b^fl/fl^*mice (AKpt) (Fig. 1A). Mice lacking the *Acvr1b^flox^* alleles were used as controls (*Ptf1a^CreER^;Kras^LSL-G12D^*) (Kpt). Using a similar approach, we combined the *Sox9CreER* allele^23,24^ with the *Kras^LSL-G12D^* and *Acvr1b^flox^* conditional alleles to express Kras^G12D^ and ablate *Acvr1b* expression in postnatal ductal cells. This generated *Sox9CreER;Kras^LSL-G12D^;Acvr1b^fl/fl^*(AKsox9) mice (Fig. 1B). *Sox9CreER;Kras^LSL-G12D^* mice (Ksox9) lacking the *Acvr1b* alleles were used as controls. All mice were injected with tamoxifen at 3-4 weeks of age and then subsets of mice were subjected to MRI imaging and survival studies, while others were collected at various time points for histological analyses.

**Figure 1.**
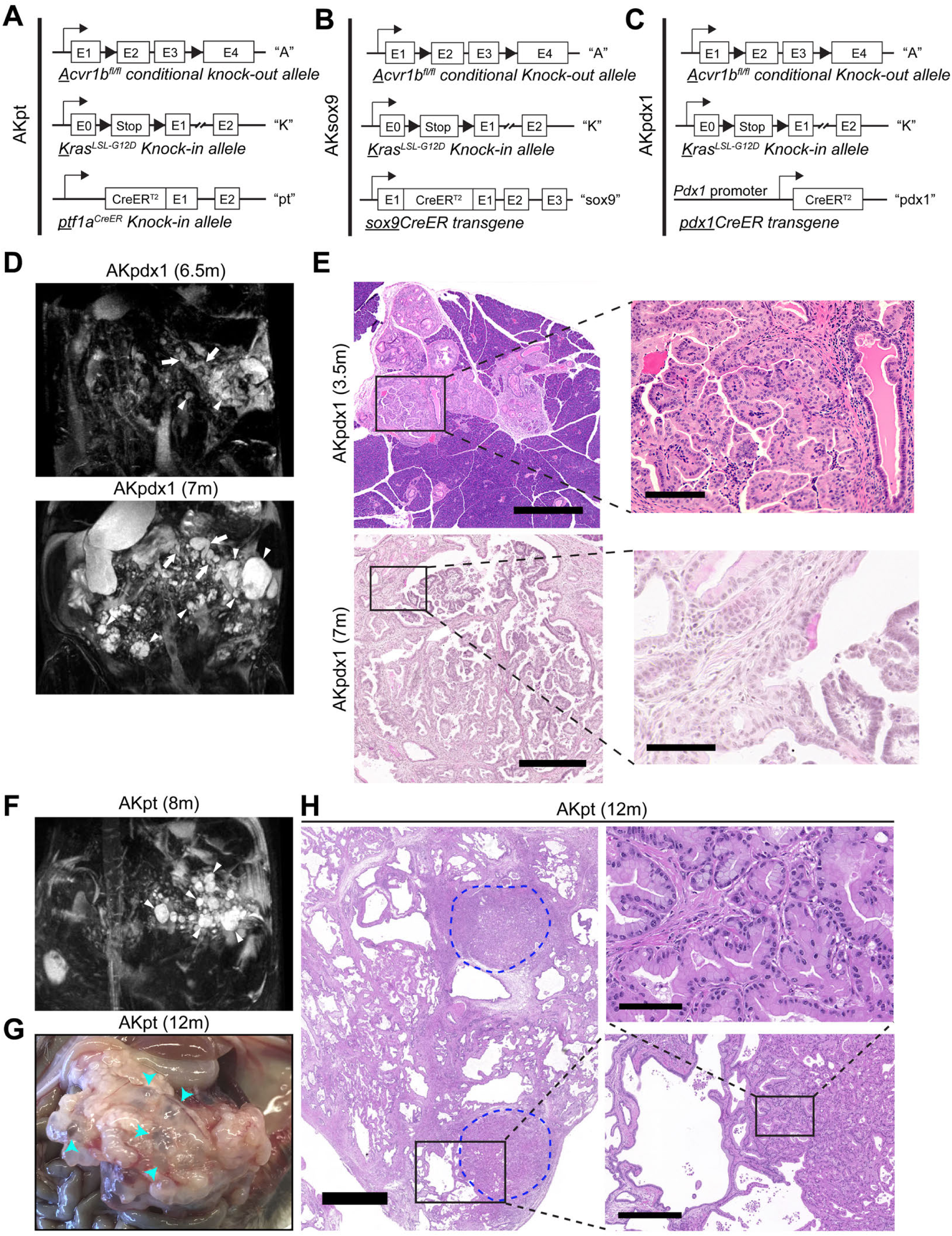
*Acvr1b* loss in Kras^G12D^-expressing acinar cells results in pancreatic cysts throughout the pancreas. **(A-C)** Schematic of the alleles in the *<u>Pt</u>f1a^CreER^;<u>K</u>ras^LSL-G12D^;<u>A</u>cvr1b^fl/fl^* (AKpt) mice **(A),** *<u>Sox9</u>CreER;<u>K</u>ras^LSL-G12D^;<u>A</u>cvr1b^fl/fl^* (AKsox9) mice (**B**), and *Pdx1Cre;Kras^LSL-G12D^;Acvr1b^fl/fl^*(AKpdx1) mice **(C)** used in this study. **(D)** Representative T2-weighted MRI images of the AKpdx1 mice (n=4) at the age of 6.5 and 7 months (m) depicting multiple cystic lesions (arrowheads), as well as dilation of the main pancreatic duct (arrows). **(E)** Low (left) and high (right) magnification images of H&E-stained sections from the pancreas of AKpdx1 mice at 3.5 or 7 months of age. **(F)** Representative T2-weighted MRI images of AKpt mice (n=3) at the age of 8 months depicting multiple cystic lesions (arrowheads), no main pancreatic duct dilation was observed. **(G)** Representative gross anatomical photograph of the abdomen from AKpt mice (n=13) at ∼12 months of age (m). Some of the numerous scattered pancreatic cysts are denoted with blue arrowheads. **(H)** H&E-stained section of the AKpt pancreas at 12 months of age showing multiple nodules of pancreatic cancer (outlined in blue and magnified in accompanying panels) amongst pancreatic cysts. Scale bars: 3 mm (H, left); 400 μm (E, left and H, right bottom); 100 μm (H, right top); 80 μm (E, right).

Survival studies with AKpt mice (n=20) and AKsox9 mice (n=5) found that AKpt mice had a median survival of 363 days (Table S1). This survival time is shorter than the reported median survival of *Pdx1Cre;Kras^G12D^*mice (∼400 days),^25^ suggesting that *Acrv1b* loss in Kras^G12D^-expressing acinar cells does reduce survival time in AKpt mice. Unlike AKpt mice, none of the AKsox9 mice reached humane endpoint by 17 months of age, which is consistent with previous studies showing that very few precancerous lesions are induced by oncogenic Kras^G12D^ expression in ductal cells.^1,2^

Our previous studies with AKpdx1 mice suggested that cystic lesions consistent with IPMN could form,^20^ therefore we also monitored a subset of AKpt (n=3) and AKsox9 (n=11) mice with MRI and compared them to AKpdx1 mice (Fig. 1C, n=4). As expected, MRI showed that all AKpdx1 mice had multiple cysts (Fig. 1D, arrowheads), as well as main pancreatic duct dilation (Fig. 1D, arrows and Table S2). Consistent with our previous studies,^20^ histological analyses of these AKpdx1 mice confirmed the presence of complex arborizing papillary lesions affecting the main pancreatic duct (Fig. 1E). In AKpt mice, two of three mice had multiple cysts detected by MRI (arrowheads, Fig 1F), but no main pancreatic duct dilation was observed in these two cases by 8 months of age (Fig. 1F and Table S2). Histological analyses of these mice confirmed the presence of large cysts lined by a morphologically normal but inflamed ductal epithelium or mucinous flat epithelium (Fig. S1). Importantly, in most AKpt mice at survival endpoint, almost the entire parenchyma was replaced with a mix of these cysts, PanIN, and other evidence of chronic pancreatitis (Fig. 1G-H). There was also pancreatic cancer in a number of AKpt mice at humane endpoint (blue outlines, Fig. 1H). In sum, AKpt mice did form MRI-detectable mucinous cysts consistent with IPMN; however, the epithelium of these cysts was primarily flat and did not recapitulate the complex arborizing papilla that could be observed in AKpdx1 mice (Fig. 1E).

MRI imaging of AKsox9 mice found that 4 of 11 mice imaged developed detectable cysts and one of the 17-month-old mice had a large cyst accompanied by clear dilation of the main pancreatic duct (Fig. 2A and Table S2). By necropsy and histologically, many of the pancreata at 12 and 17 months of age were predominantly normal (Fig. 2B) and the cysts that were present were predominantly lined by flat, reactive ductal cells (Fig. 2C), however, some mice also had mucinous PanIN lesions (Fig. 2D). Together these results indicated that *Acvr1b* loss and activation of Kras in ductal cells can result in ductal cysts that affect the main pancreatic duct. However, similar to AKpt mice, the papillary nodules present in AKpdx1 mice were not present in AKsox9 mice.

**Figure 2.**
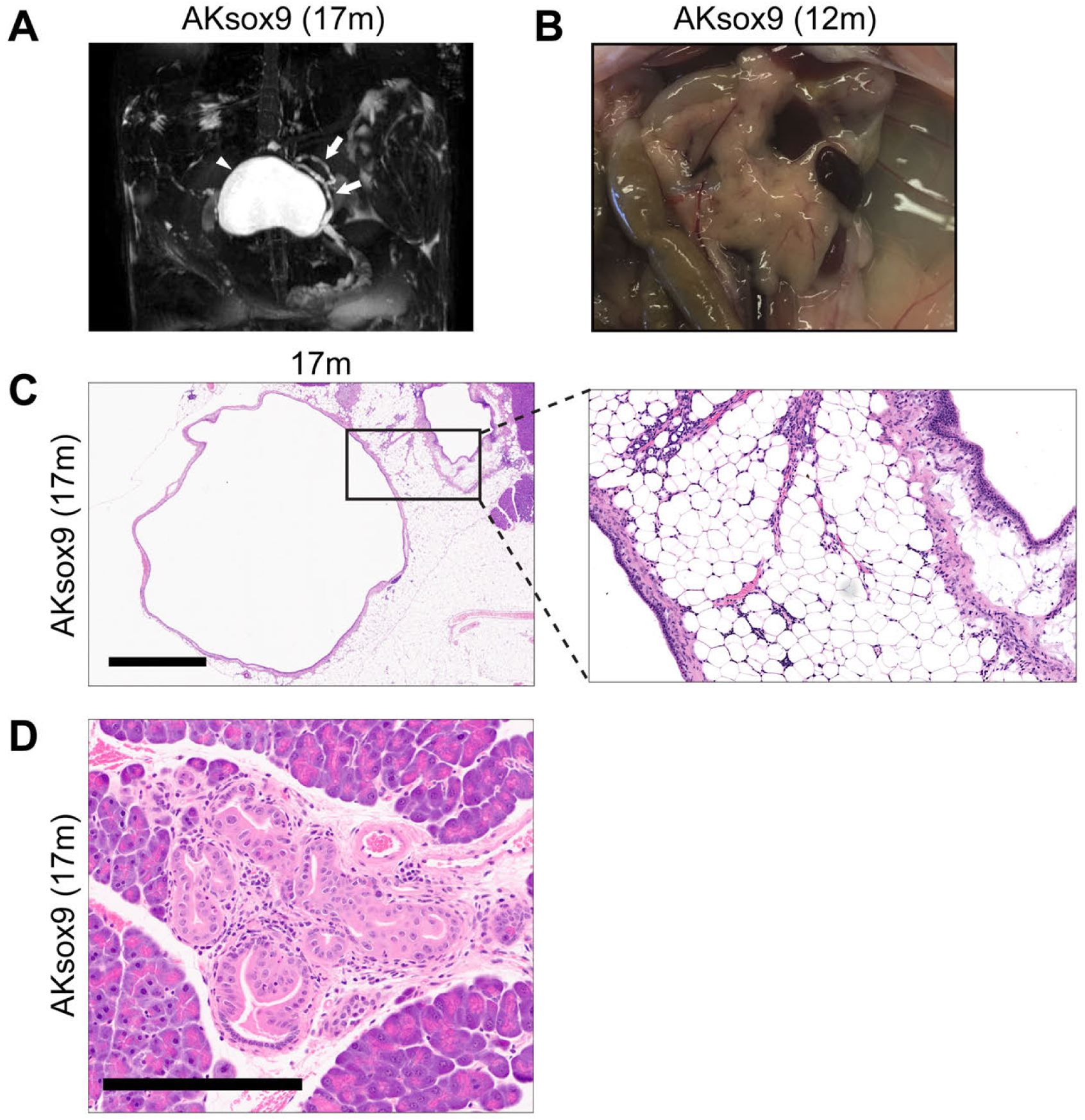
*Acvr1b* loss and activation of Kras in ductal cells is associated with dilation of the main pancreatic duct. **(A)** Representative MRI images of *Sox9CreER;Kras^LSL-G12D^;Acvr1b^fl/fl^*(AKsox9) mice (n=5) at 17 months (m) of age depicting a large cyst (arrowhead), as well dilation of the main pancreatic duct (arrows). **(B)** Representative gross anatomical photographs of the mouse abdomen from AKsox9 mouse at 12 months of age. (n=13) **(C-D)** H&E-stained sections of AKsox9 pancreata at 17 months of age depicting a large non-mucinous cyst and dilation of main pancreatic duct beside it (C, left) and fatty fibrosis between the two large ducts (C, right). **(D)** Example of a PanIN lesion found in AKsox9 mice at 17 months of age. Scale bars: 1.2 mm (C, left); 200 μm (D).

### *Acvr1b* loss and activation of Kras in acinar cells promotes formation of PanIN and IPMN-like precancerous lesions

To further characterize the events leading to cyst formation in AKpt mice, we histologically characterized AKpt mice at 3, 5, 7, and 9 months of age (Fig. 3A). At 3 months of age, small areas of the AKpt pancreata had ductal metaplasia that replaced normal acinar cell area (acinar-to-ductal metaplasia (ADM)), as well as low-grade PanIN lesions (Fig. 3A). In 5-month-old AKpt mice, ADM and immune cell infiltration was more prevalent (Fig. 3A). In addition to low-grade PanIN, high grade PanIN were also observed and these became more prevalent at 7 months of age (Fig. 3A). In 9-month-old AKpt mice, large areas of the pancreas were displaced by large ductal cysts with reactive epithelium or mucinous characteristics (Fig. 3A). One of three 7- and 9-month-old AKpt mice also developed PDAC (Fig. 3A). In contrast, in 7, 9, and 12-month-old Kpt mice (Fig. 3B), more normal parenchyma remained and the histological changes consisted mostly of ADM or low-grade PanIN lesions, with very rare large cystic ducts. These results suggested that the combined effects of Kras mutation and *Acvr1b* loss in acinar cells resulted in formation of more precancerous lesions compared to Kras activation alone.

**Figure 3.**
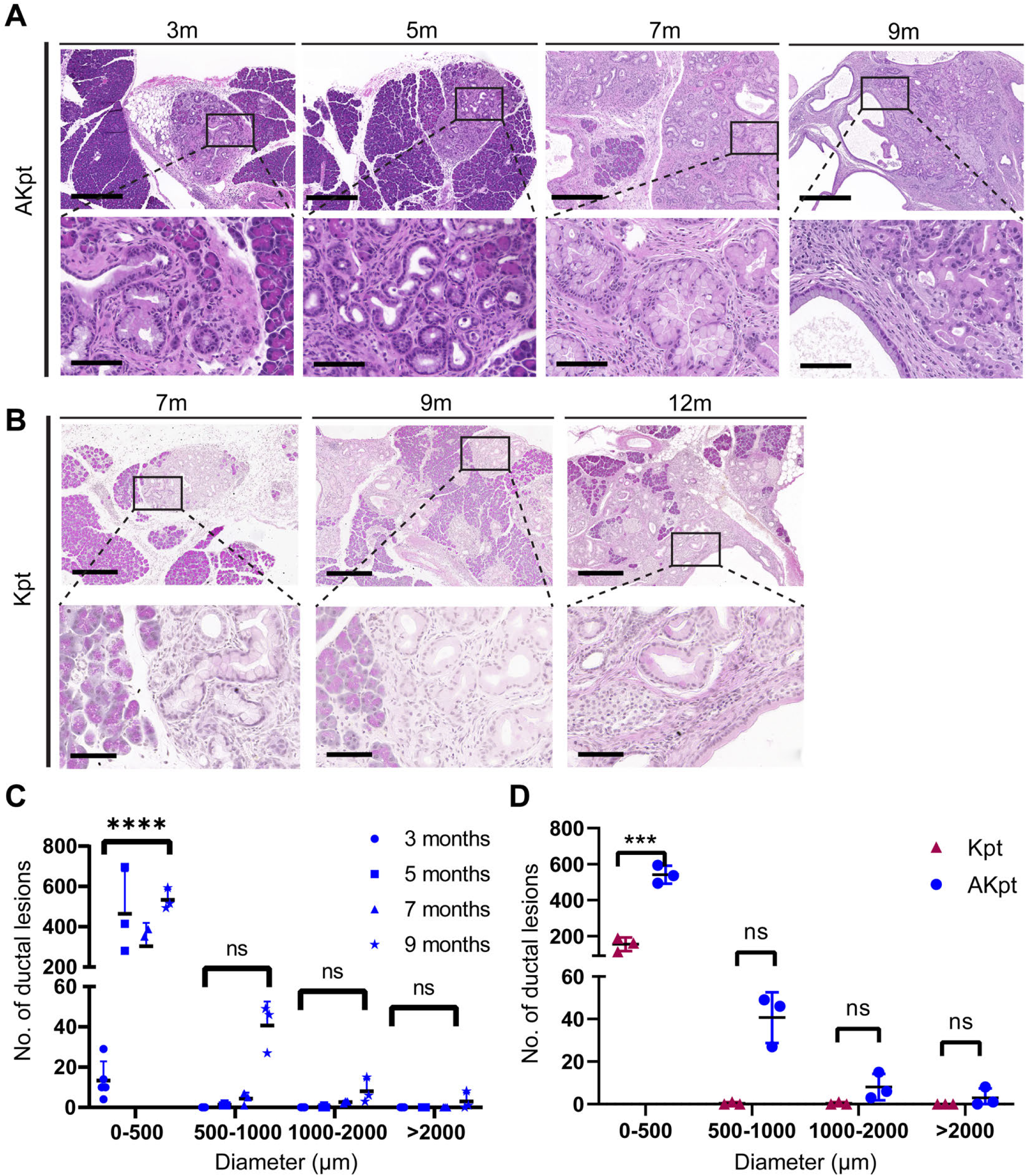
*Acvr1b* loss and activation of Kras in acinar cells promotes initiation of pancreatic cysts with preneoplastic characteristics. **(A-B)** H&E-stained sections of *<u>Pt</u>f1a^CreER^;<u>K</u>ras^LSL-G12D^;<u>A</u>cvr1b^fl/fl^* (AKpt) pancreata at 3 (n=5), 5 (n=3), 7 (n=3), and 9 months of age (n=3) **(A)** or *<u>Pt</u>f1a^CreER^;<u>K</u>ras^LSL-G12D^* (Kpt) mice at 7, 9, and 12 months of age (n=3) **(B)**. **(C-D)** Graphs depicting the number of the neoplastic structures at different size groups: 0-500 μm, 500-1000 μm, 1000-2000 μm and >2000 μm in AKpt mice at 3 (n=5), 5 (n=3), 7 (n=3), and 9 months of age (n=3) **(C)** or in Kpt (n=3) vs. AKpt (n=3) mice at the age of 9 months **(D)**. P-values: **** < 0.0001, *** <0.001. Scale bars: upper image of set, 400 μm; lower image of set, 80 μm (A-B).

To more accurately quantify the histological changes in the ductal lumens across AKpt pancreata collected at different ages (3, 5, 7, and 9 months), we measured the diameter and subcategorized abnormal ductal lesions into four size groups: 0-500 µm, 500-1000 µm, 1000-2000 µm and >2000 µm (Fig. 3C and Table S3). At 3 months of age, all the ductal lesions were smaller than 500 μm (Fig. 3C). However, from 5 months of age onwards, we observed ductal structures larger than 500 μm, with an increase in the overall number of structures as the AKpt mice aged (Fig. 3C). Additionally, we observed some abnormal ductal structures larger than 2000 μm at the age of 9 months (Fig. 3C). Altogether, these findings suggested that the AKpt pancreata progressively become more cystic and this coincides with the progressive loss of normal parenchyma and wide-spread histological evidence of chronic pancreatitis (Fig. 3A). We further compared the number and size of ductal structures in AKpt (n=3) and Kpt mice (n=3) at the age of 9 months (Fig. 3D and Table S4). We found that AKpt mice had more ductal structures larger than 500 μm when compared to Kpt mice, as well as significantly more ductal lesions less than 500 μm (Fig. 3D). These data suggested that loss of *Acvr1b* in Kras^G12D^-expressing acinar cells results in increased preneoplastic formation overall and large cysts reminiscent of flat gastric IPMN-like lesions (>500 μm) form at later ages. Additionally, this increased lesion formation is accompanied by evidence of chronic pancreatitis.

### *Acvr1b* loss and activation of Kras also promotes induction of precancerous lesions from ductal cells

To further characterize the events associated with cyst formation in AKsox9 mice, we histologically characterized the pancreata of AKsox9 and Ksox9 mice at 5, 7, 9, or 12 months of age (Fig. 4A and Fig. S2A) and measured the diameter of the preneoplastic lesions in AKsox9 mice at 5, 7, 12, and 17 months of age (Fig. 4B and Table S5). Consistent with previous studies,^2^ few PanIN lesions were observed in Ksox9 mice and only two of six mice examined had any PanIN at 5 or 9 months of age (Fig. S2A and Table S5). At 5 months of age, the AKsox9 pancreata were normal and no PanIN or pancreatitis was observed (Fig. 4A and Table S5). At 7, 12, and 17 months of age, preneoplastic lesions were observed in AKsox9 mice and the number of lesions per mouse increased with age (Fig. 4A-B). These preneoplastic lesions were predominantly classified as PanIN, however, there were also larger mucinous cysts consistent with main duct IPMN of the flat, gastric type (Fig. 4C and Table S5). This suggests that loss of *Acvr1b* in ductal cells increases the formation of preneoplastic lesions from Kras^G12D^-expressing ductal cells.

**Figure 4.**
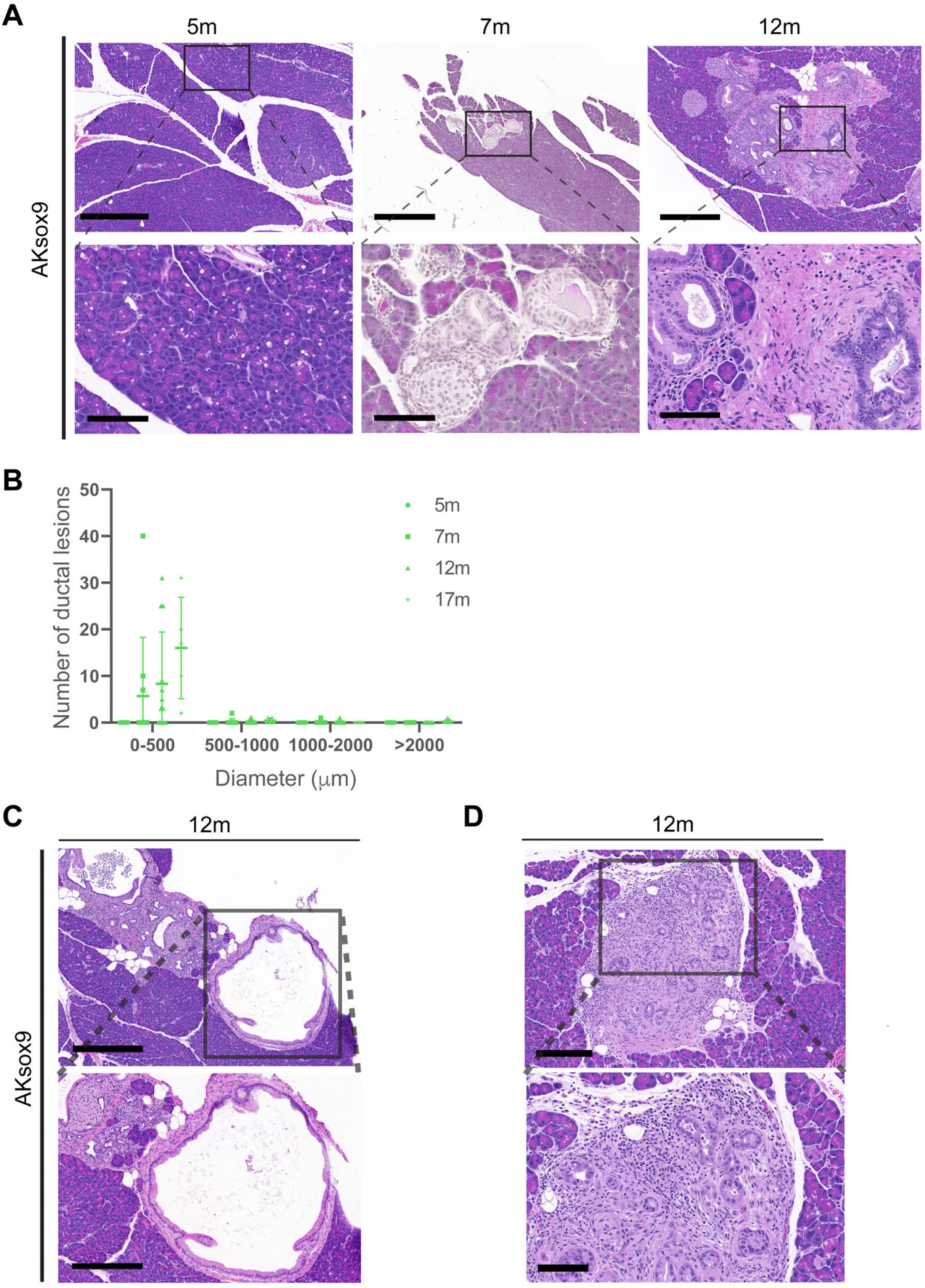
*Acvr1b* loss and activation of Kras in ductal cells promotes formation of preneoplastic lesions and cysts. **(A)** Representative images from H&E-stained sections of the *Sox9CreER;Kras^LSL-G12D^;Acvr1b^fl/fl^* (AKsox9) pancreata at 5 (n= 5), 7 (n= 10), and 12 months of age (n= 13). **(B)** Graph depicting the number of neoplastic structures in different size groups: 0-500 μm, 500-1000 μm, 1000-2000 μm, and >2000 μm in the AKsox9 mice at 5 (n=5), 9 (n=10), 12 (n=13), and 17 months of age (n=5). **(C-D)** H&E-stained sections from AKsox9 mice at 12 months of age depicting a large preneoplastic cyst **(C)** and chronic pancreatitis **(D)**. Scale bars: 480 μm (C, upper image); 300 μm (C, lower image); 200 μm (D, upper image, 100 μm (A, D, lower image).

In addition to the observed precancerous lesions, we noted that there was evidence of chronic pancreatitis in multiple AKsox9 mice, including the formation of nodules of inflamed ducts, large cysts with reactive ductal epithelium, and fatty fibrosis surrounding the remnants of islets and ducts (Fig. 2C, 4A, 4D, and Table S5). This suggests that similar to *Pdx1Cre;Acvr1b^fl/fl^*mice,^20^ loss of *Acvr1b* in ductal cells, at least in the context of KrasG12D expression, is associated with the development of chronic pancreatitis as the mice age.

### Preneoplastic lesions induced by *Acvr1b* loss and Kras activation have a gastric phenotype

Low-grade PanIN lesions and MCN lesions have a predominantly gastric type epithelium, while IPMN can be divided into four main epithelial phenotypes: gastric, intestinal, pancreatobiliary, and oncocytic type. Discrimination of these phenotypes is largely based on size, histological, and morphological features, as well as the immunohistochemical expression pattern of MUC1, MUC2 and MUC5AC.^26–34^ Because none of the lesions in AKpdx1, AKpt, or AKsox9 mice possessed the highly cellular stroma associated with MCN lesions (Fig. 1-4),^35^ we focused here only on the histological and immunohistochemical pattern of mucin expression. Histopathological examination of IPMN-like lesions in AKpdx1 (3-month-old), AKpt (12-month-old), and AKsox9 (12-month-old) mice demonstrated that a small number of the cysts in AKpdx1 mice had a prominent proliferation of epithelial cells comprised of complex papillary projections (Fig. 5). The papillary epithelium showed various grades of dysplasia from low grade to high grade. The preneoplastic cells in these AKpdx1 mice were composed of flat and papillary epithelium with abundant mucinous cytoplasm and were positive for Alcian blue staining (Fig. 5). Immunohistochemical staining for MUC1, MUC2, and MUC5AC (Fig. 5) showed that these lesions expressed MUC1 and MUC5AC, but there was no expression of MUC2 (Fig. 5). The mucinous IPMN-like cysts in AKpt and AKsox9 mice lacked the complex papillary projections and had a flat gastric epithelium (Fig. 5). Consistent with this morphology, they were also positive for MUC1, MUC5AC, and Alcian Blue, but negative for MUC2. In sum, these findings indicate that the IPMN-like lesions in AKpdx1 mice are consistent with either gastric or pancreatobiliary IPMN, while those in the AKpt and AKsox9 mice have the features of gastric IPMN.

**Figure 5.**
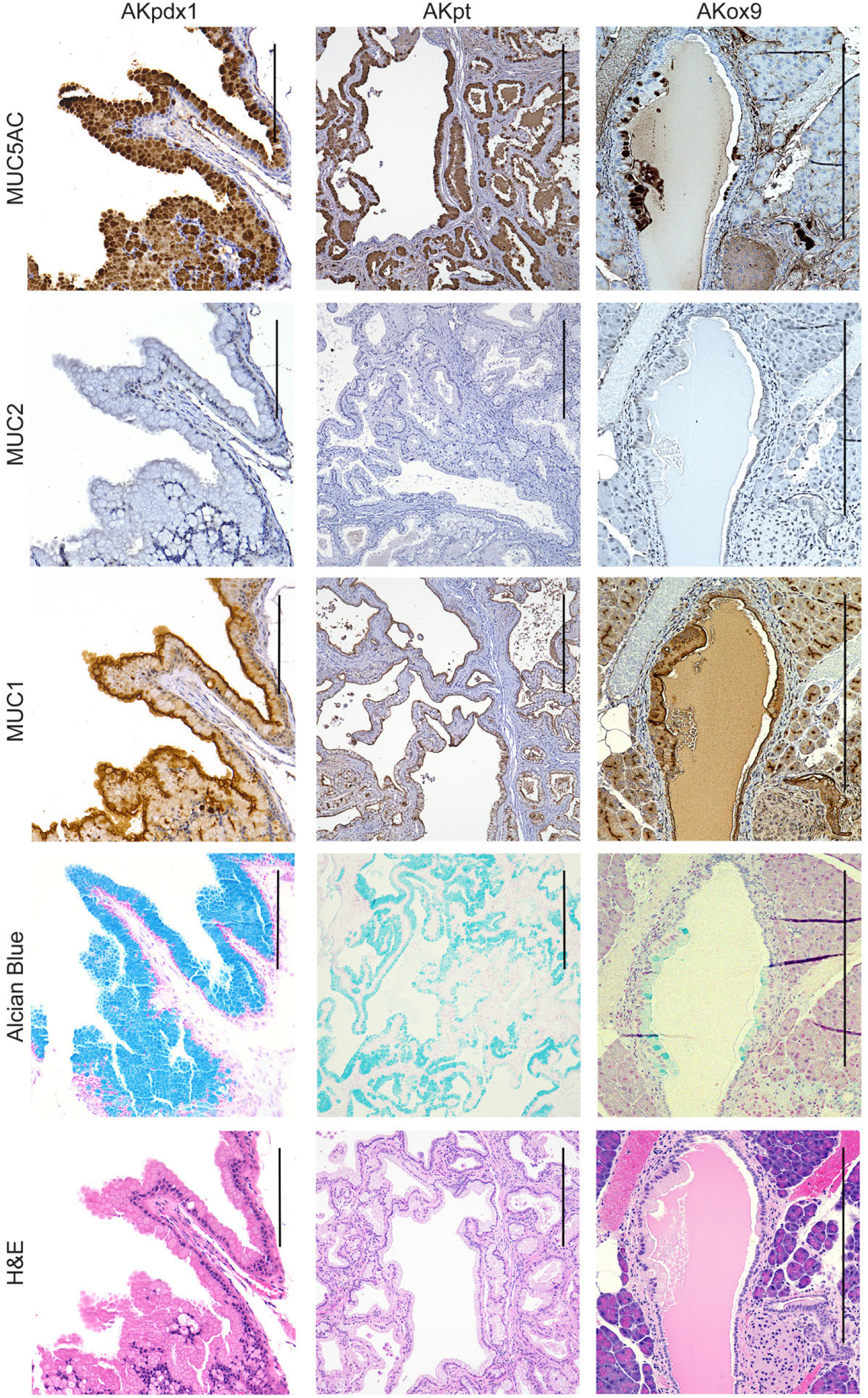
Morphological and immunohistochemical analysis of the preneoplastic lesions in AKpdx1, Akpt, and AKsox9 mice. Representative images of H&E, Alcian blue, MUC1, MUC2 and MUC5AC staining of the pancreas from mucinous large preneoplastic cysts in *Pdx1Cre;Kras^LSL-G12D^;Acvr1b^fl/fl^*(AKpdx1), *Ptf1a^CreER^;Kras^LSL-G12D^;Acvr1b^fl/fl^*(AKpt), or *Sox9CreER;Kras^LSL-G12D^;Acvr1b^fl/fl^* (AKsox9) mice. Scale bars: 200 μm.

### Caerulein treatment accelerates formation of IPMN-like lesions in AKpt mice

Pancreatitis is known to accelerate the development of ADM/PanIN in the presence of oncogenic Kras and it is a risk factor for pancreatic cancer.^2,36–39^ Since chronic pancreatitis was prominent in older AKpt mice and this was coincident with cyst formation, we next examined whether experimentally induced acute pancreatitis would also be sufficient to promote increased cyst formation in AKpt mice. Acute pancreatitis can be experimentally induced by treatment with supraphysiological levels of the cholecystokinin receptor agonist, caerulein.^40^ To evaluate the impact of acute pancreatitis on the formation of PanIN and IPMN-like lesions following acinar cell-specific Kras activation with or without *Acvr1b* loss, we injected tamoxifen at 3-4 weeks of age and then induced caerulein-mediated acute pancreatitis in 6-week-old Kpt (n=4) and AKpt mice (n=4) and analyzed pancreatic tissue 21 days later (Fig. 6A-B). Consistent with previous literature,^2^ we confirmed that acute pancreatitis resulted in PanIN formation from oncogenic Kras-expressing acinar cells in Kpt mice by day 21 (Fig. 6B). In the AKpt mice, acute pancreatitis increased the replacement of acinar cells with ductal-like cells compared to Kpt mice, and more lesions with larger ductal lumens were observed (Fig. 6B and Fig. S3). To quantify the number of acinar cells in AKpt and Kpt mice, we performed IHC to detect amylase in the pancreas of the caerulein-treated Kpt (n=4) and AKpt mice (n=4) and measured the amylase positive pancreatic areas. The percentage of amylase-positive area in the AKpt mice was significantly lower than that of the Kpt mice (p<0.05) (Fig. 6C and Table S6). We also performed immunohistochemical staining for CK19 on Kpt (n=4) and AKpt pancreata (n=4) at day 21 and quantified the CK19-positive areas. We found that the proportion of CK19-positive area in the AKpt mice was significantly higher than that of the Kpt mice (p<0.05) (Fig. 6D and Table S6). Together these data indicate that loss of the activin receptor in AKpt mice resulted in a greater proportion of the parenchyma being affected 21 days after the induction of pancreatitis. We observed no difference in the relative amounts of CK19-positive ADM and PanIN lesions between genotypes, however the epithelial lumens tended to be bigger on average with a larger number of IPMN-like cysts (lumen size >500 μm) in AKpt compared to Kpt mice (Fig. 6E and Fig. S3). In light of the absence of IPMN-like lesions in 3-month-old AKpt mice (Fig. 3C, Table S3), these results suggest that pancreatitis contributes to cyst development AKpt mice.

**Figure 6.**
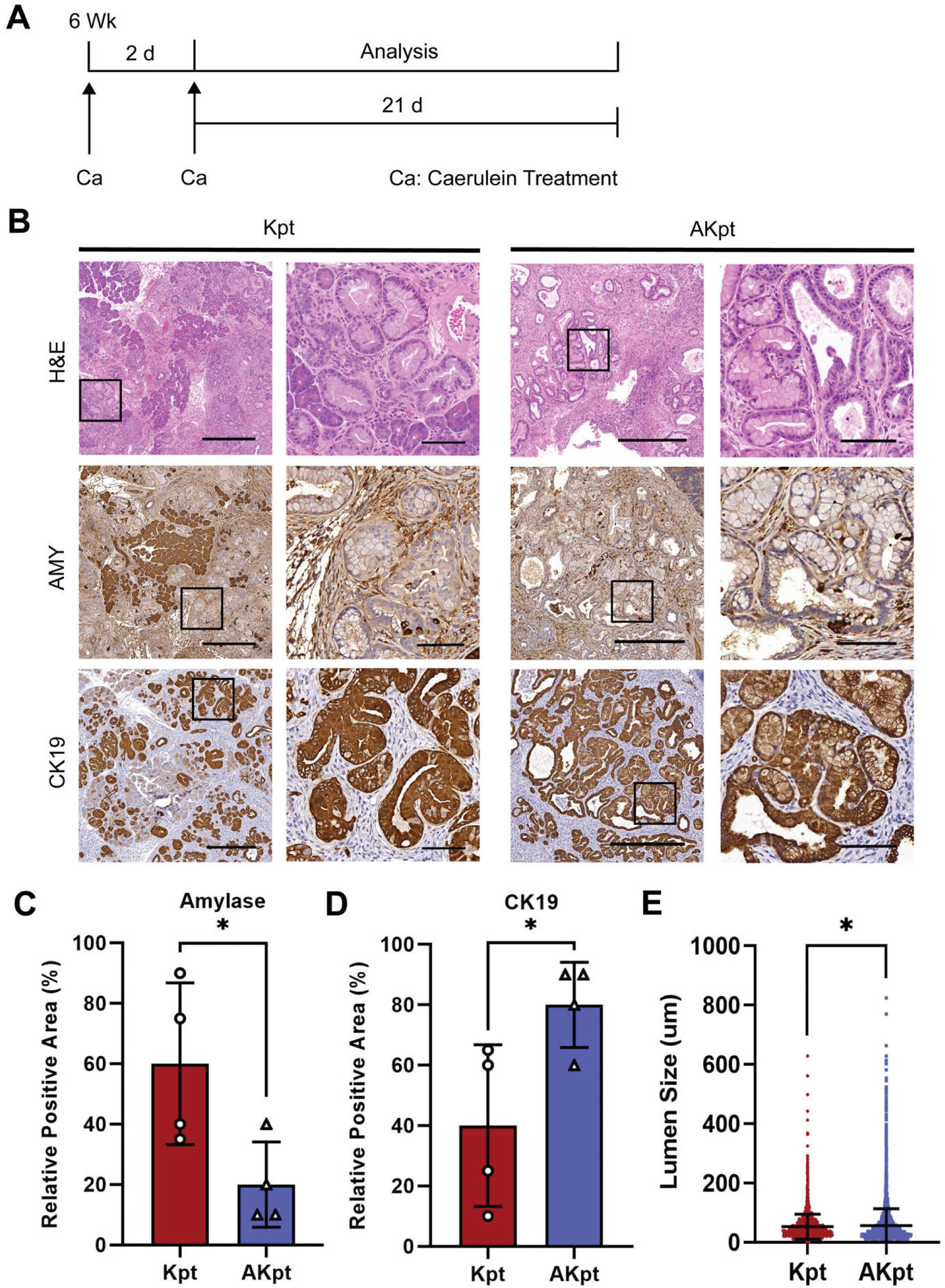
Caerulein treatment and *Acvr1b* loss promoted formation of IPMN-like lesions from Kras^G12D^-expressing acinar cells. **(A)** Schematic of the caerulein treatment plan - At 6 weeks of age, *Ptf1a^CreER^;Kras^LSL-G12D^*(Kpt) (n=4) and *Ptf1a^CreER^;Kras^LSL-G12D^;Acvr1b^fl/fl^*(AKpt) (n=4) mice were treated with two rounds of caerulein (Ca) injections on alternative days and analyzed 21 days later. **(B)** Representative images of H&E staining of the pancreata from Kpt and AKpt mice treated with caerulein at day 21. Immunohistochemistry staining against Amylase (AMY) or CK19 in Kpt or AKpt pancreata 21 days after caerulein treatment. **(C-D)** Quantification of the Amylase-positive **(C)** or CK19-positive **(D)** pancreatic areas in the Kpt mice compared to the AKpt mice at 21 days after the caerulein treatment. **(E)** Quantification of lumen size across the entire section in Kpt or AKpt mice 21 days after caerulein treatment. P-values: * < 0.05 by t-test **(C-D)** and non-parametric t-test **(E)**. Scale bars: (B) left image of each set, 400 μm; right image of each set, 80 μm.

### Precancerous lesions are not induced by caerulein treatment of AKsox9 mice

IPMN and PanIN formation was also coincident with evidence of chronic pancreatitis in AKsox9 mice, therefore, we next examined how caerulein-mediated acute pancreatitis affects PanIN and IPMN formation from Kras^G12D^-expressing ductal cells with or without *Acvr1b* loss. To do this, we injected tamoxifen at 3-4 weeks of age and then induced caerulein-mediated acute pancreatitis in 6-week-old Ksox9 (n= 4-6) and AKsox9 mice (n= 3-4) and analyzed pancreatic tissue 2 or 21 days and 9 weeks later (Fig. 7A). Similar amounts of inflammation and degranulation of the acinar cells was observed at 2 days after caerulein treatment in both genotypes, but we did not detect any PanINs in either Ksox9 or AKsox9 mice 21 days or 9 weeks after caerulein treatment (Fig. 7B). These results suggested that short-term acute pancreatitis did not promote Kras^G12D^-expressing ductal cells to form preneoplastic lesions or cysts regardless of the *Acvr1b* genotype.

**Figure 7.**
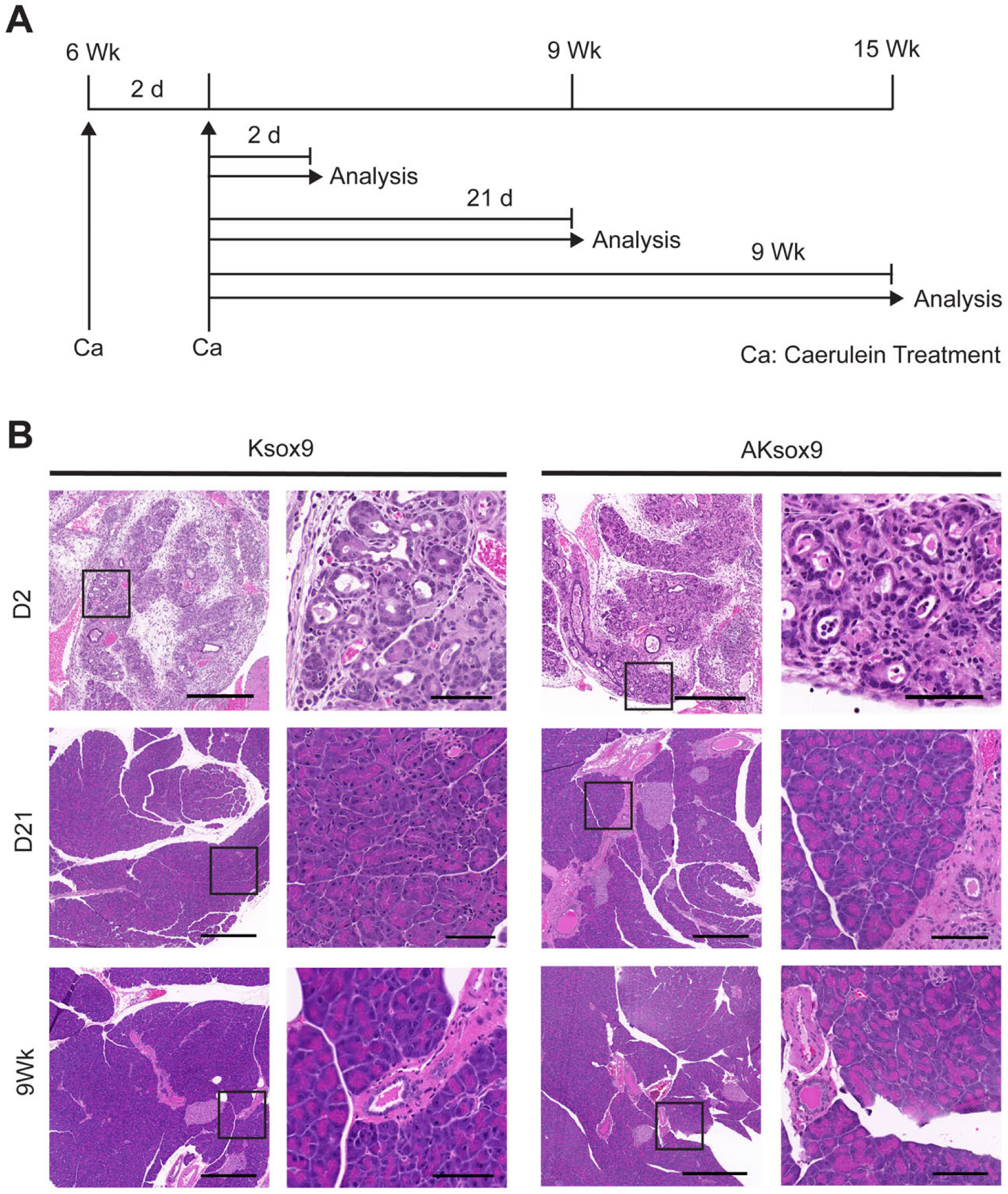
Kras expressing ductal cells have a low propensity to form PanINs after chemically-induced acute pancreatitis. **(A)** Schematic of the caerulein treatment plan - At 6 weeks of age, *Sox9CreER;Kras^LSL-G12D^;Acvr1b^fl/fl^* (AKsox9) (n= 4-6) and *Sox9CreER;Kras^LSL-G12D^* (Ksox9) (n= 3-4) mice were treated with two rounds of caerulein (Ca) injections on alternative days and analyzed at day 2 or 21 (D), or 9 weeks (wk) later. **(B)** Representative images of H&E staining of the pancreas from Ksox9 or AKsox9 mice treated with caerulein, 2 or 21 days or 9 weeks post treatment. Scale bars: (B) left image of each set, 400 μm; right image of each set, 80 μm.

## Discussion

### In the context of oncogenic Kras, embryonic loss of *Acvr1b* is not equivalent to loss of *Acvr1b* postnatally

Our previous studies suggested that IPMN resembling the gastric or pancreatobiliary subtypes were induced by oncogenic Kras and loss of *Acvr1b* in the pancreas.^20^ Here, in this study, we found that inducing these same mutations in just the postnatal ductal or acinar cell populations did not result in IPMN with the complex arborizing papillae of pancreatobiliary IPMN. However, flat, mucinous, and large-lumened ductal cysts, consistent with gastric type IPMN, as well as smaller lumen sized PanIN, were observed, which is similar to the predominant phenotype found in AKpdx1 mice.^20^ There are a number of possible reasons why the phenotype in AKpdx1 may not be recapitulated in the AKpt or AKsox9 mice. First, our previous studies suggested that loss of *Acvr1b* alone during pancreatogenesis did not affect pancreatic development because the minor pancreatitis phenotype was not observed until after 8 months of age.^20^ However, AKpdx1 mice formed cysts as early as 1-month-old;^20^ this suggests that oncogenic Kras and loss of *Acvr1b* in combination might have affected some aspect of pancreatic development or the postnatal maturation of the pancreas and resulted in cyst, and thus IPMN, formation. Second, many pancreatic cell types are targeted when the *Pdx1Cre* allele is used. This can result in compound effects from both acinar and ductal cells being affected. Consistent with this possibility, the combined loss of *Acvr1b* and Kras activation was associated with chronic pancreatitis in AKsox9 mice (Fig. 4D). Therefore, it is possible that pancreatitis that occurs as a result of loss of *Acvr1b* in Kras^G12D^-expressing ductal cells in AKpdx1 mice, could have further exacerbated the phenotype of Kras^G12D^-expressing, *Acvr1b*-deleted acinar cells and vice versa. Lastly, as opposed to the AKsox9 and AKpt mouse models, islet cell types are also affected in AKpdx1 mice. Endocrine cells of the islet, particularly beta cells, are known to affect initiation of precancerous lesions by insulin/insulin receptor signaling in acinar cells.^41–43^ Since Activin also affects insulin secretion from islets,^44,45^ it is plausible that changes in Activin signaling in multiple cell types could result in a different phenotype in AKpdx1 mice compared to AKsox9 or AKpt mice.

### *Acvr1b* loss and activation of Kras in either the acinar or ductal cell lineages promotes formation of precancerous lesions

Our study demonstrates that loss of *Acvr1b* in juvenile ductal or acinar cells expressing Kras^G12D^ results in the formation of more precancerous lesions compared to Kras^G12D^ expression alone. In addition, some of the lesions in AKpt and AKsox9 mice become large cystic lesions with flat mucinous epithelium, consistent with gastric type IPMN. To date, the majority of animal models of pancreatic cancer have focused on examining the effect of cancer associated mutations in all pancreatic cells or by specifically examining the effect in acinar cells.^6^ Thus, knowledge of how genetic changes affect tumorigenesis are largely based on the effects on all pancreatic cells in the context of an embryonic induction or on juvenile or adult acinar cells. Relatively recent studies have begun to identify how pancreatic-cancer-associated genetic changes affect ductal cells’ contribution to pancreatic cancer *in situ*. The effects of the mutations studied, thus far, have shown that some of the mutations that affect tumorigenesis from ductal cells have little or opposite effects on acinar cells.^46–48^ In our studies, we found that *Acvr1b* loss, likely through abrogation of Activin signaling, is one of the few pathways identified to date that promotes precancerous lesion formation from both acinar and ductal cells expressing Kras^G12D^. However, the molecular mechanism by which this occurs is still unknown. It appears that rather than directly promoting precancerous lesion formation, Activin/Acvr1b signaling might affect the extent of inflammation in the pancreas that subsequently promotes formation of precancerous lesions. Previous studies have found that the expression of Activin increases in the epithelium during injury and Activin then acts on the macrophages to activate them.^49^ Additionally, Activin-blocking antibody treatment of mice injected with caerulein reduced the severity of caerulein-induced pancreatitis in wild-type mice.^49^ Thus, Activin signaling is clearly involved in pancreatitis, but further studies of ductal- or acinar-cell specific Activin/Acvr1b signaling are needed to further understand how loss of *Acvr1b* affects the inflammatory signaling milieu in the pancreas in the presence or absence of Kras mutations.

### Genetic changes in acinar and ductal cells can contribute to formation of IPMN-like lesions

Our data suggests that loss of *Acvr1b* in Kras^G12D^-expressing ductal or acinar cells can lead to formation of large mucinous lesions akin to IPMN lesions, as well as cysts lined with non-mucinous epithelium. Previous studies have predominantly reported that only ductal cells can give rise to IPMN-like lesions, while the same mutations in acinar cells typically induce PanIN lesions or have no effect.^2,48,50,51^ Rare instances of IPMN-like lesions occurring in the context of acinar-cell specific mutations have been observed^52,53^. Indeed, Flowers et al. used tdTomato lineage labeling with the same *Ptf1aCreER* allele used in our study to show that IPMN can be derived from Ptf1a^+^ cells.^53^ In our studies, it appears that effects of *Acvr1b* loss on cyst formation is not limited to either acinar or ductal cells; but due to the increased effects of Kras^G12D^ on the acinar cell population likely leads to a more consistent and robust phenotype in AKpt mice compared to AKsox9 mice. Specifically, in AKpt mice, more and/or earlier precancerous lesions form compared to Kpt mice. As a result, ductal metaplasia replaces the majority of the normal parenchyma and the ongoing inflammation from this metaplasia is associated with the formation of large non-mucinous and mucinous cysts, like IPMN. This suggests that in the AKpt model inflammation is, in part, important for the cystic phenotype. This is likely similar to role that *Elastase*-driven TGFalpha expression has on IPMN formation in the context of Kras^G12D^ expression in the entire pancreas.^52^ In sum, experiments in mice suggest that it is possible for acinar cells to contribute to IPMN, but the applicability of these observations to human IPMN would be strengthened by future studies examining whether IPMN can originate from acinar cells when only a few of these cells sustain IPMN-associated mutations so that the large effects that occur after loss of the entire parenchymal function can be separated from the actions of the mutations in IPMN induction from acinar cells.

In AKsox9 mice, *Acvr1b-*loss*-*associated inflammation is also likely involved in the formation of main-duct IPMN and non-mucinous cysts. However, because *Kras^G12D^* mutations alone have very little effect on ductal cells,^1,2,54^ this likely resulted in fewer precancerous lesions when compared to AKpt mice. Additionally, the large effects on the parenchyma in AKpt mice did not occur in AKsox9 mice. This makes interpreting the relationship between the inflammation and the induction precancerous lesions more straightforward. For example, it is easier to identify a mucinous lesion obstructing the main duct and conclude that it is downstream from the dilation and inflammation observed in an upstream area of the pancreas in these mouse models. In sum, inflammation likely has a large role in producing the conditions leading to formation of large cystic lesions that can be classified as IPMN from ductal cells. This duct-centric inflammation as a causative agent in IPMN formation seems consistent with location of main duct IPMN primarily within the pre-existing ducts in patients.

## Methods

### Mouse Strains

*LSL-Kras^G12D^*,^21^ *Acvr1b^fl/fl^*,^20^ *Ptf1a^CreER^*(Jax No. 019378),^22^ and *Sox9CreER*^23,24,46^ mice have been described in detail previously. To induce recombination, 3-4-week-old mice were administered three subcutaneous injections of tamoxifen (Sigma Aldrich) in corn oil over 5 days (days 1, 3, and 5) at 5 mg/40 g body weight. *Ptf1a^CreER^;Kras^LSL-G12D^*(Kpt) mice and *Sox9CreER;Kras^LSL-G12D^* (Ksox9) mice were used as controls for *Ptf1a^CreER^;Kras^LSL-G12D^;Acvr1b^fl/fl^* (AKpt) mice and *Sox9CreER;Kras^LSL-G12D^;Acvr1b^fl/fl^*(AKsox9) mice. No developmental effects were observed in these mice. All mice were housed in the Animal Care Facility at Columbia University Irving Medical Center (CUIMC), and the studies were conducted in compliance with the CUIMC Institutional Animal Care and Use Committee (IACUC) guidelines. Specifically, rapid weight loss in adjunct with body condition score less than 3 (Ullman-Cullere et al, Laboratory Animal Science, 1999), debilitating diarrhea, progressive dermatitis, rough hair coat, hunched posture, lethargy or persistent recumbency, coughing, labored breathing, nasal discharge, neurologic signs, bleeding from any orifice, self-induced trauma, shivering, a tumor burden of >10% body weight, ulceration of the tumor or a mean tumor diameter >20 mm were considered a humane end point. If the animals lost more than 20% of their body weight, were cachectic, unable to obtain food or water, or their ocular and/or respiratory systems were compromised, they were euthanized.

### Histological and Immunolabeling Analyses

Murine tissues were fixed overnight in 10% neutral buffered formalin and embedded in paraffin. Routine H&E staining was performed by the Histology Service Core Facility at CUIMC. K.S. and I.W. evaluated the histology of all sections and J.L.K, D.F.S, G.H.S., and Dr. Wanglong Qiu (University Irvine Medical Center) provided a secondary analysis of key sections and all figures.

For histological analyses, one section per mouse was examined in its entirety and the number of ductal lesions/lumens and their respective diameters were measured.

Immunohistochemistry was performed using unstained 5-μm sections derived from the formalin-fixed and paraffin-embedded blocks that were deparaffinized in xylene 3 times and rehydrated in ethanol 4 times. Heat-induced antigen retrieval was performed on all slides in Tris-EDTA buffer (0.5% Tween-20, citrate buffer, pH 6.8) in a steamer for 30 minutes. Slides were incubated with Dako peroxidase block buffer to block endogenous peroxidase activity. Primary antibody staining was performed at 4°C overnight. The secondary antibodies were detected with a 40-minute incubation with biotinylated universal secondary antibodies (Dako, Carpinteria, CA) and streptavidin–horseradish peroxidase. Hematoxylin was then used as a counterstain. Slides were dehydrated in ethanol then xylene and mounted with VectaMount permanent mounting medium (Vector Laboratories Burlingame, CA). The following primary antibodies were used for immunohistochemistry: Amylase (1:100, cat: sc-46657, Santa Cruz), CK19 (1:500, cat: ab52625, Abcam), MUC1 (1:100, #BS-1497R, Bioss Inc.), MUC2 (SC-15334, Santa Cruz) and MUC5AC (#BS-7166R, Bioss Inc.).

For quantification of the IHC (CK19 and Amylase), one section per mouse was examined in its entirety and the percentages of the CK19 or Amylase positive areas in each mouse were calculated by their proportions to the entire section.

For Alcian blue staining, 5-μm sections were deparaffinized and rehydrated as in immunohistochemistry staining. Then slides were stained with Alcian blue solution (1 g Alcian blue in 100 ml 3% glacial acetic acid, pH 2.5) for 30 minutes at room temperature. Counterstaining with 0.1% nuclear fast red solution (0.1 g nuclear fast red and 5 g ammonium sulfate in 100 ml dH2O) was performed for 5 minutes. Slides were dehydrated and mounted.

Images were primarily captured using a 3D Histotech slide scanner and exported using the Aperio ImageScope program.

### Caerulein treatment

Acute pancreatitis was induced at 6 weeks of age in AKpt, Kpt, AKsox9, and Ksox9 mice by 2 sets of 6 hourly i.p. caerulein injection (50 μg/kg diluted in saline; Sigma-Aldrich) separated by 24 hours. In this experiment, the final day of the caerulein/saline injection was considered day 0^2,36^ and the pancreata were collected as indicated.

### MRI experiment

MRI data were obtained on a Bruker BioSpec 9.4T Magnetic Resonance Imager (Bruker Corp., Billerica, MA). The mice were anesthetized with 1-2% isofluorane mixed with medical air via a nose cone. The concentration of the isoflurane was adjusted during the procedure to maintain the respiration rate in the range of 40-70 breaths/min using a respiration pillow attached to a monitoring system (SA Instruments, Stony Brook, NY). Body temperature was maintained around 37^°^C using a flowing water heating pad. Low-resolution T1 weighted scout images were obtained initially. A rapid acquisition with relaxation enhancement sequence was used to acquire a high-resolution T2 weighted images with the following parameters: repetition time (TR) = 4873 ms, echo time (TE) = 60 ms, field of view = 34 × 34 mm, matrix size =256 x 256, slice thickness = 0.4 mm with 30 slices to cover the entire abdomen. Maximum Intensity Projection (MIP) technique was used for the visualization of the cysts.

### Statistical Analysis

*P* values were calculated using the 2-way ANOVA with GraphPad Prism (version 8.0.2). P values <0.05 were determined to be significant.

## Acknowledgements

The authors thank members from Su and Kopp labs for discussions. We thank Christopher Wright (Vanderbilt, USA) for the *Ptf1a*^CreER^ mice and Maike Sander for the *Sox9CreER* mice. We thank the Histology Service Core Facility at CUIMC.

---

**Figure S1.**
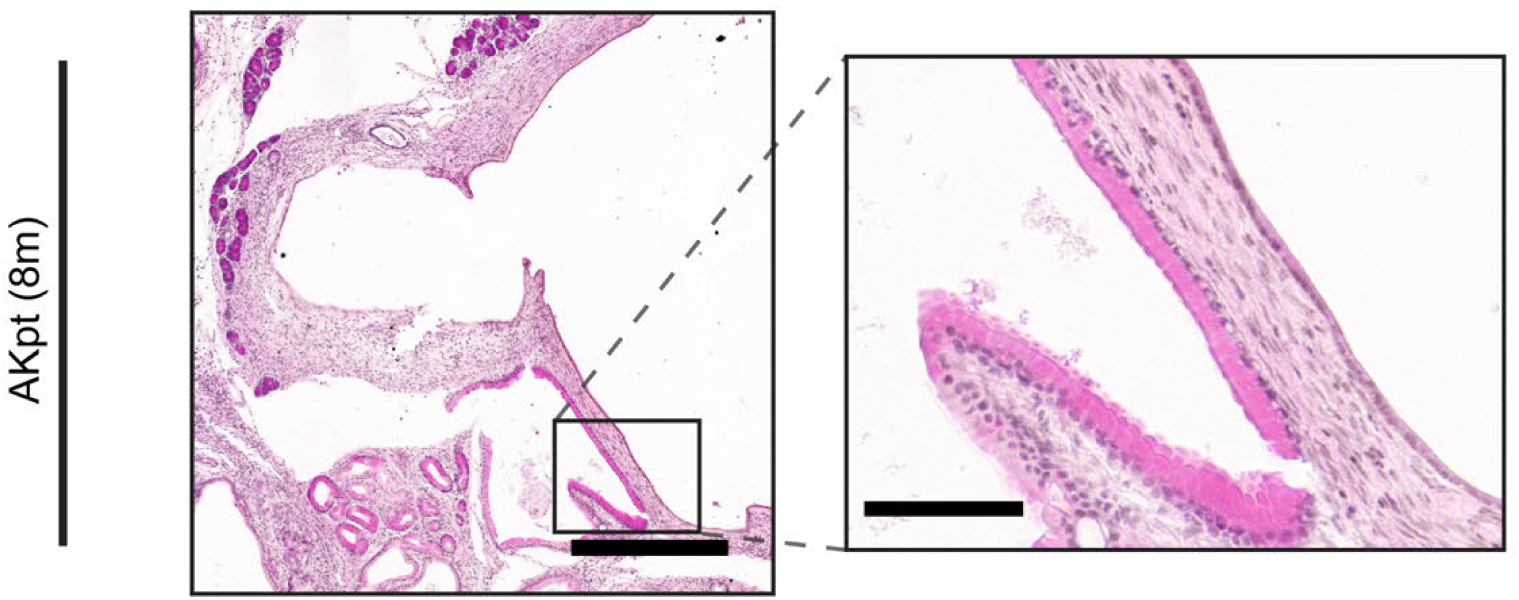
Histological images corresponding to MRI imaged AKpt pancreata. Low and high magnification images of H&E-stained sections from *Ptf1a^CreER^;Kras^LSL-G12D^;Acvr1b^fl/fl^* (AKpt) pancreata at 8 months of age. Scale bars: left image, 400 μm; right image, 80 μm.

**Figure S2.**
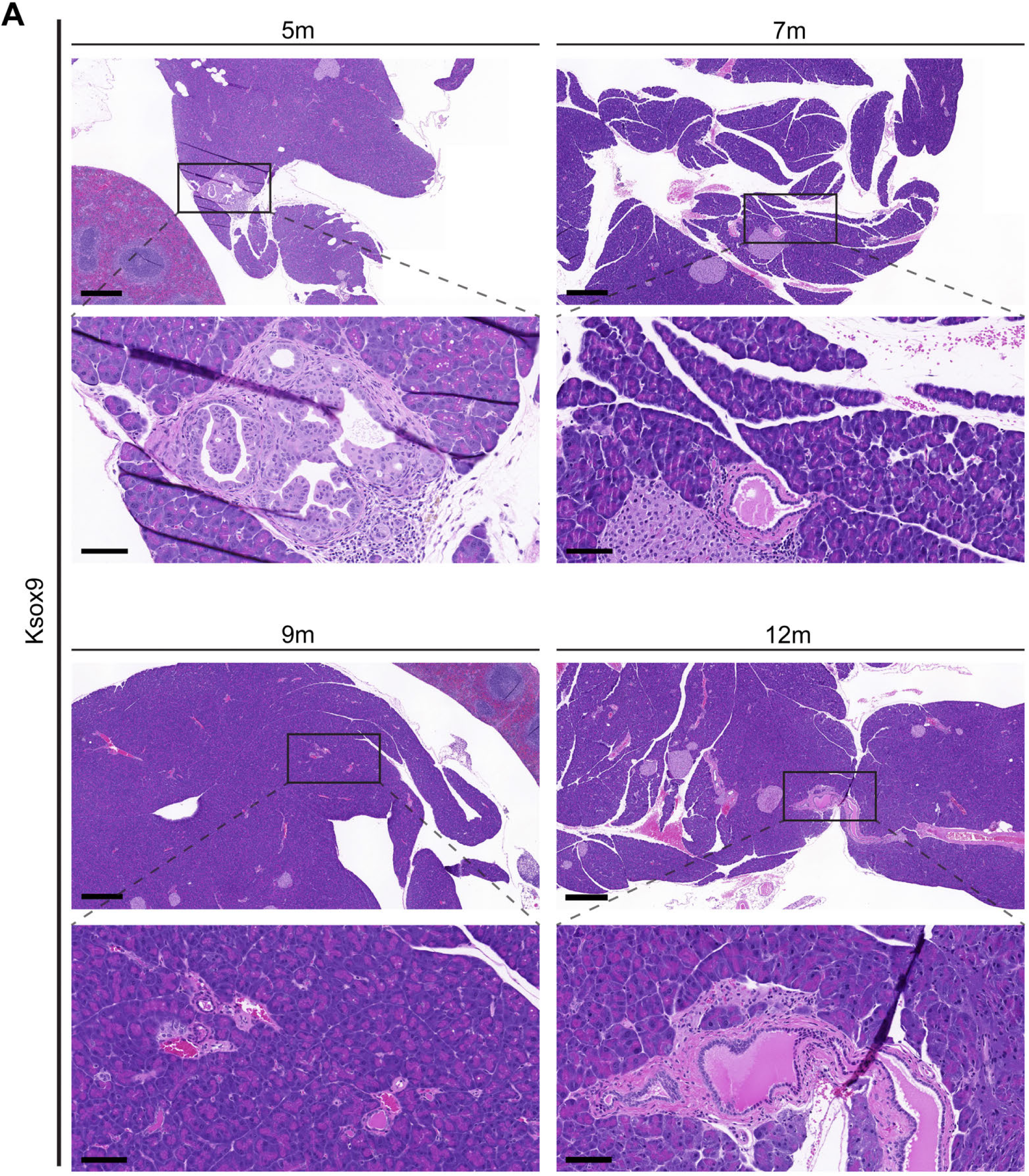
Ductal-cell-specific expression of oncogenic Kras results in few PanINs lesions. H&E-stained sections of the pancreas from *Sox9CreER;Kras^LSL-G12D^*(Ksox9) mice at 5, 7, 9, or 12 months of age. Out of the six 5- and 9-month-old Ksox9 mice analyzed only one mouse at 5 months old and one mouse at 9 months old had PanINs (Table S4). Scale bars: upper image per set, 400 μm; lower image per set, 80 μm.

**Figure S3.**
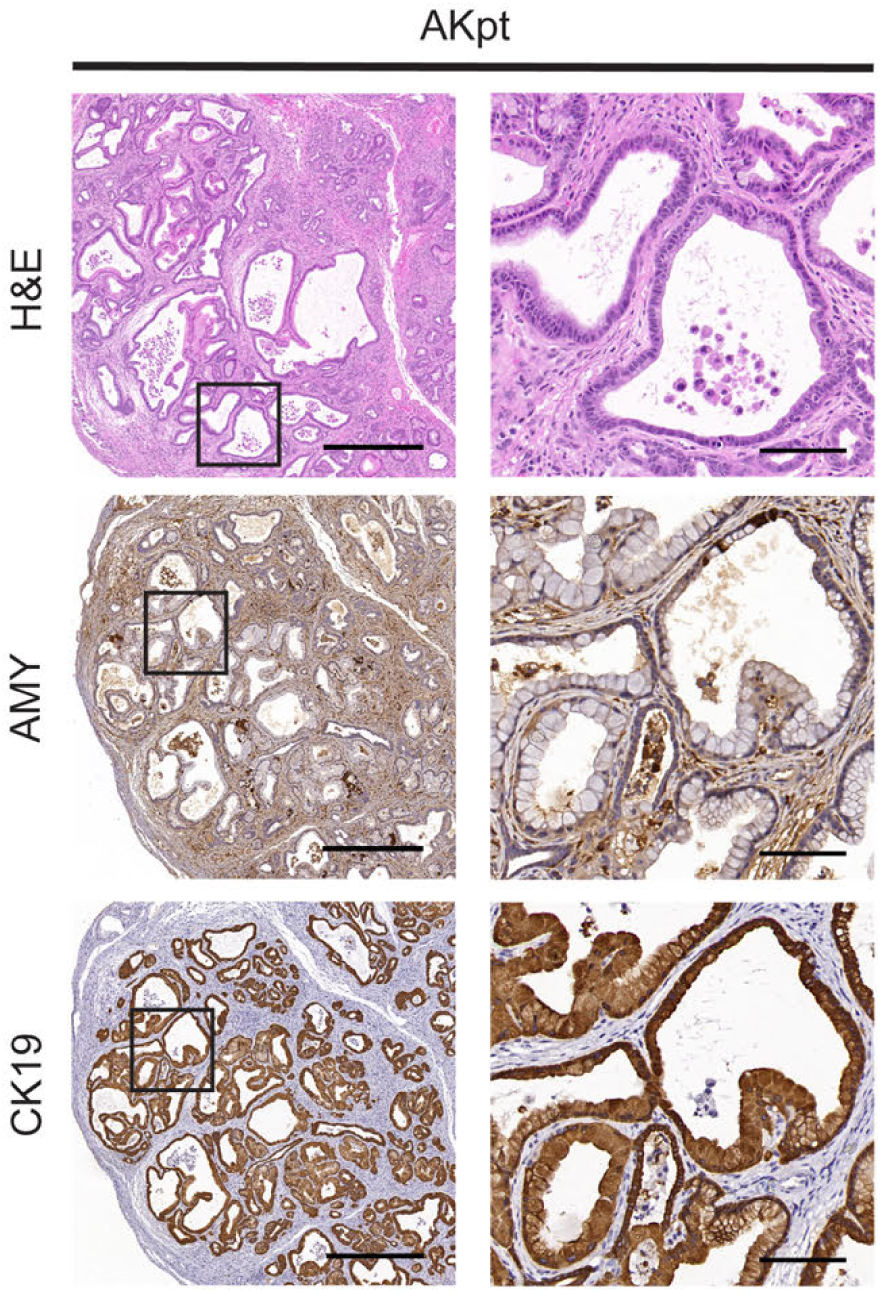
Caerulein treatment promoted formation of IPMN-like lesions in AKpt mice. Representative images of H&E staining of the pancreata from *Ptf1a^CreER^;Kras^LSL-G12D^* (Kpt) and *Ptf1a^CreER^;Kras^LSL-G12D^;Acvr1b^fl/fl^*(AKpt) mice treated with caerulein. Immunohistochemistry staining against Amylase (AMY) or CK19 in Kpt or AKpt pancreata 21 days after caerulein treatment. Scale bars: left image of each set, 400 μm; right image of each set, 80 μm.

